# Variation in assembly stoichiometry in non-metazoan homologs of the hub domain of Ca2+/Calmodulin-dependent protein kinase II

**DOI:** 10.1101/536094

**Authors:** Ethan D. McSpadden, Zijie Xia, Chris C. Chi, Anna C. Susa, Neel H. Shah, Christine L. Gee, Evan R. Williams, John Kuriyan

**Author notes:** To whom correspondence should be addressed: John Kuriyan, Kuriyan Lab, 176 Stanley Hall, MC 3220, University of California, Berkeley, CA 94720-3220, USA. Present address: Intel corporation, Santa Clara, CA, USA. Present address: Department of Chemistry, Columbia University, New York City, NY, USA.

## Abstract

The multi-subunit Ca^2+^/calmodulin-dependent protein kinase II (CaMKII) holoenzyme plays a critical role in animal learning and memory. The kinase domain of CaMKII is connected by a flexible linker to a C-terminal hub domain that assembles into a 12- or 14-subunit scaffold that displays the kinase domains around it. Studies on CaMKII suggest that the stoichiometry and dynamic assembly/disassembly of hub oligomers may be important for CaMKII regulation. Although CaMKII is a metazoan protein, genes encoding predicted CaMKII-like hub domains, without associated kinase domains, are found in the genomes of some green plants and bacteria. We show that the hub domains encoded by three related green algae, *Chlamydomonas reinhardtii*, *Volvox carteri f. nagarensis*, and *Gonium pectoral*, assemble into 16-, 18-, and 20-subunit oligomers, as assayed by native protein mass spectrometry. These are the largest known CaMKII hub domain assemblies. A crystal structure of the hub domain from *Chlamydomonas reinhardtii* reveals an 18-subunit organization. We identified four intra-subunit hydrogen bonds in the core of the fold that are present in the *Chlamydomonas* hub domain, but not in metazoan hubs. When six point mutations designed to recapitulate these hydrogen bonds were introduced into the human CaMKII-α hub domain, the mutant protein formed assemblies with 14 and 16 subunits, instead of the normal 12- and 14-subunit assemblies. Our results show that the stoichiometric balance of CaMKII hub assemblies can be shifted readily by small changes in sequence.

## Introduction

CaMKII (Ca^2+^/calmodulin-dependent protein kinase II) is a multimeric Ser/Thr kinase that plays critical roles in neuronal signaling and cardiac pacemaking in animals ^1–4^. Each molecule of CaMKII consists of a kinase domain and a calmodulin-binding regulatory segment that are connected by a variable-length linker to a hub domain, also referred to as the association domain (**Figure 1A**). Hub domains oligomerize into donut-shaped assemblies of 12 or 14 subunits ^5–7^, leading to holoenzymes in which the kinase domains are arranged around the assembled hub ^8,9^ (**Figure 1B**). This multimeric architecture underlies many of the remarkable biochemical properties exhibited by CaMKII, including an ultrasensitive response to Ca^2+^/CaM ^8,10–12^.

**Figure 1.**
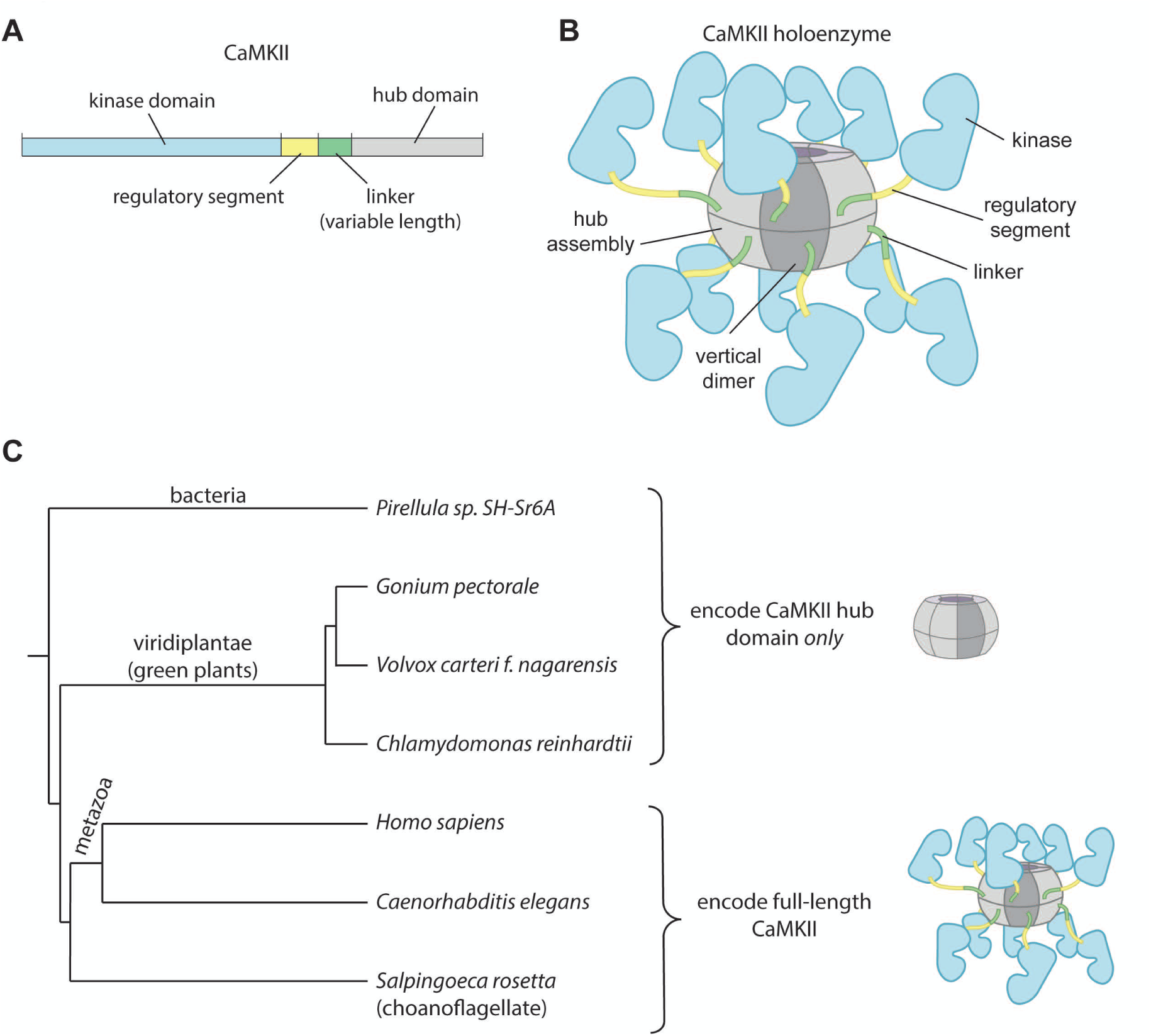
(**A**) Domain organization for an intact CaMKII subunit. (**B**) Depiction of the CaMKII holoenzyme, C-terminal hub domains (shown in gray) oligomerize into a donut shaped structure, forming the core of the holoenzyme. N-terminal kinase domains (shown in blue) extend outwards from the core, tethered to the hub by a flexible linker. The holoenzyme depicted here is a dodecamer, but tetradecameric holoenzymes can be formed as well (Bhattacharyya et al. 2016; Myers et al. 2017). In this diagram, which represents the activated state of CamKII, the autoinhibitory regulatory segments (yellow) are shown displaced from the kinase domains. (**C**) A phylogenetic tree of some organisms that encode forms of CaMKII. Full length CaMKII is only found in metazoans and premetazoan choanoflagellates. Isolated CaMKII hub domains are found in the genomes of some green algae and some bacteria.

The CaMKII hub domain is related structurally to a diverse group of proteins that include nuclear transport factor 2 (NTF2) and ketosteroid isomerases, but it shares no significant sequence similarity with these proteins ^5^. Like the proteins to which it is related structurally, the CaMKII hub domain forms a stable wedge-shaped dimer ^7^. The dimers of the CaMKII hub oligomerize to form the ring-shaped structures that hold the holoenzyme intact (we refer to these dimers as “vertical dimers,” because they span the equatorial plane of the hub assembly: see Figure 1B). The linker connecting the kinase domain to the hub varies in length between the four CaMKII isoforms in humans (denoted the α, β, γ, and δ isoforms).

CaMKII holoenzymes undergo activation-triggered destabilization and subunit exchange ^7,13^. Although the mechanism of this phenomenon is not completely understood, it suggests that the number of subunits in a hub assembly may not be static. Crystal structures of mammalian CaMKII hubs have revealed both tetradecameric forms (composed of seven vertical dimers, PDB codes 1HKX ^5^ and 2W2C ^12^) and dodecameric forms (six vertical dimers, PDB codes 5IG3 ^7^ 3SOA ^8^ and 2UX0 ^12^). Electron microscopic images of negatively-stained samples of intact, full-length human CaMKII-α show both dodecameric and tetradecameric forms ^7,9^. The oligomerization state of the mouse CaMKII-α hub has been observed to change from dodecameric to tetradecameric forms in response to the removal of the kinase domains, either through gene truncation or proteolytic cleavage ^6^.

Homologs of CaMKII have been found so far only in animals and choanoflagellates, the closest extant relatives of the true metazoans ^7^. The sequences of the kinase domain and the hub domain are highly conserved across mammalian and choanoflagellate species ^7,14^. Crystal structures have been determined for the hubs of CaMKII variants from two invertebrate species, the worm *C. elegans* and the sea anemone *N. vectensis*. Both form tetradecameric assemblies (*C. elegans* PDB code 2F86 ^6^, *N. vectensis* PDB code 5IG4 ^7^). A different CaMKII hub isoform from *N. vectensis* crystallized as a cracked, partially open tetradecamer (PDB code 5IG5 ^7^). The CaMKII hub from a choanoflagellate, *S. rosetta*, forms an open spiral assembly rather than a closed ring (PDB code 5IG0 ^7^).

Given that full-length CaMKII proteins are restricted to metazoans and the choanoflagellates, it is surprising that genes with roughly 50% sequence identity to the human CaMKII-α hub domain are present in the genomes of several green algae and bacteria (**Figure 1C**). These organisms do not appear to contain sequences homologous to the CaMKII kinase domain. We report here the biophysical characterization of several of these non-metazoan CaMKII hub domains, with a particular emphasis on the protein from the green algae *Chlamydomonas reinhardtii*. The crystal structure of the *Chlamydomonas* hub assembly reveals that the structure of individual hub domains within the assembly is essentially the same as seen in the human hub assembly, as expected from sequence similarity. Unexpectedly, we observed, both by crystallography and native mass spectrometry, that the *Chlamydomonas* hub assembles as an 18-subunit closed ring, notably larger than the dodecameric or tetradecameric human CaMKII hub assemblies.

Changes in the stoichiometry of hub assemblies, between dodecameric, tetradecameric, and spiral forms, are correlated with changes in the twist and curvature of the central β-sheet of the hub domain ^7^. The *Chlamydomonas* hub contains several intra-subunit hydrogen bonds that are not present in the human hub domains, and these appear to support the increased twist of the β-sheet that is necessary for forming a larger assembly. Guided by the *Chlamydamonas* hub structure, we introduced six mutations into the human CaMKII-α hub with the goal of promoting the formation of these hydrogen bonds. Under native mass spectrometry conditions the mutant human CaMKII-α hub formed assemblies with14 and 16 subunits, rather than the 12- and 14-subunit assemblies seen normally. A crystal structure of the mutant human CaMKII-α hub domain was obtained in a tetradecameric form, confirming the formation of the desired hydrogen bonds. Our results show how small changes in the sequence can shift the balance between different stoichiometries of the CaMKII hub assembly.

## Results and Discussion

### Identification of non-metazoan CaMKII hub domains

Genes with approximately 50% sequence identity to the human CaMKII-α hub domain, without associated kinase domains, were identified by a protein BLAST search in the genomes of three closely related species of green algae, *Chlamydomonas reinhardtii* (UniprotKB A8IHL6 ^15^), *Volvox carteri f. nagarensis* (UniprotKB D8U3T0 ^16^), and *Gonium pectoral* (UniprotKB A0A150GVZ6 ^17^). The sequences of these algal hub domains share approximately 80% identity with each other. Sequences with ~50% identity to the human CaMKII-α hub domain were also found in several species of bacteria, including *Pirellula sp. SH-Sr6A* (UniprotKB A0A142Y204) and *Planctopirus sp. JC280* (UniprotKB A0A1C3EH80) (**Figure 2**). None of these algal or bacterial genomes appear to contain a kinase domain related to CaMKII, either in the open reading frame corresponding to the hub domain, or elsewhere in the genome, and so the function of these isolated hub domains is unclear. We shall refer to the proteins we have identified as “CaMKII hubs,” although we stress that these non-metazoan species do not appear to contain actual CaMKII-like kinases.

**Figure 2.**
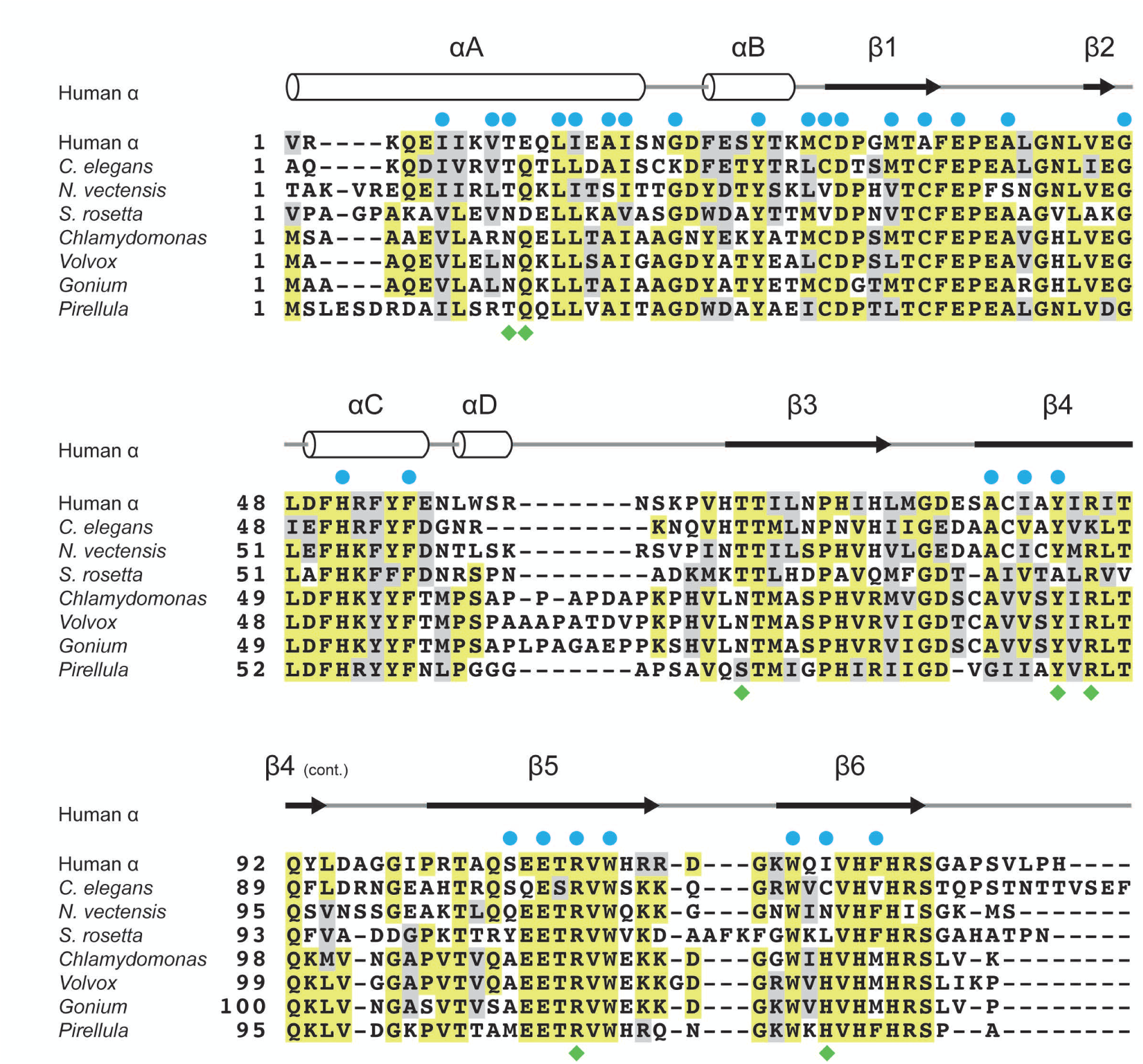
Sequence alignment of the human CaMKII-α hub domain and the hub domains encoded by *C. elegans*, *N. vectensis*, *S. rosetta*, *Chlamydomonas*, *Volvox*, *Gonium and Pirellula*. The human to *Chlamydomonas* point mutation sites that shifted the preferred oligomerization states of the human hub domain are marked with green diamonds. The residues that are buried in the *Chlamydomonas* and mutant human hub domain structures are marked with a blue circle. Yellow indicates high sequence identity and grey represents partial sequence identity.

### Oligomerization states of non-metazoan CaMKII hub domains

The three algal CaMKII hubs were expressed individually in *E. coli* and purified to homogeneity. Native protein mass spectrometry was used to determine the molecular weight of oligomers formed by the purified proteins. The *Chlamydomonas* CaMKII hub domain formed an oligomer of 309,105 ± 400 Daltons, corresponding to a complex of 18 subunits (See Table I for molecular masses and estimated subunit stoichiometry for all complexes analyzed. No other oligomeric states were observed (**Figure 3A**). The oligomerization state was found to be concentration independent over the range of 8 to 400 µM, suggesting that the detected 18-subunit complex is a stable assembly.

**Figure 3.**
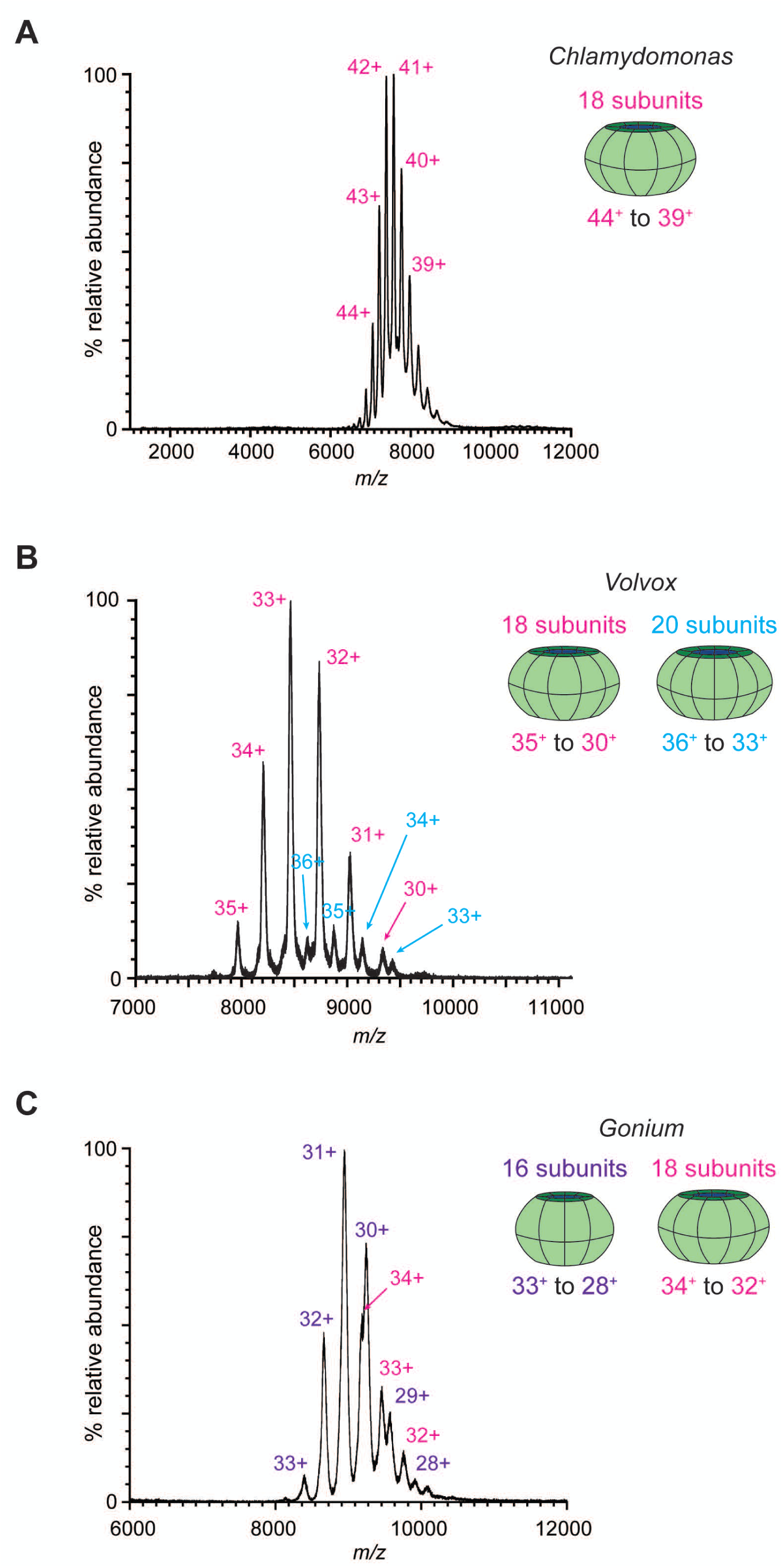
(**A**) Electrospray ionization mass spectrum (ESI-MS) under native conditions of the *Chlamydomonas* CaMKII hub domain. A single species was detected with a molecular weight of 309,105 Da, corresponding to an 18-mer. (**B**) ESI-MS of the *Volvox* CaMKII hub domain. Two species were detected with molecular weights of 275, 440 and 309,860 Da, corresponding to 18-mers and 20-mers, respectively. (**C**) ESI-MS of the *Gonium* CaMKII hub domain. Two species were detected with molecular weights of 275,440 and 309,860 Da, corresponding to 16-subunit and 18-subunit assemblies, respectively.

**Table I:**
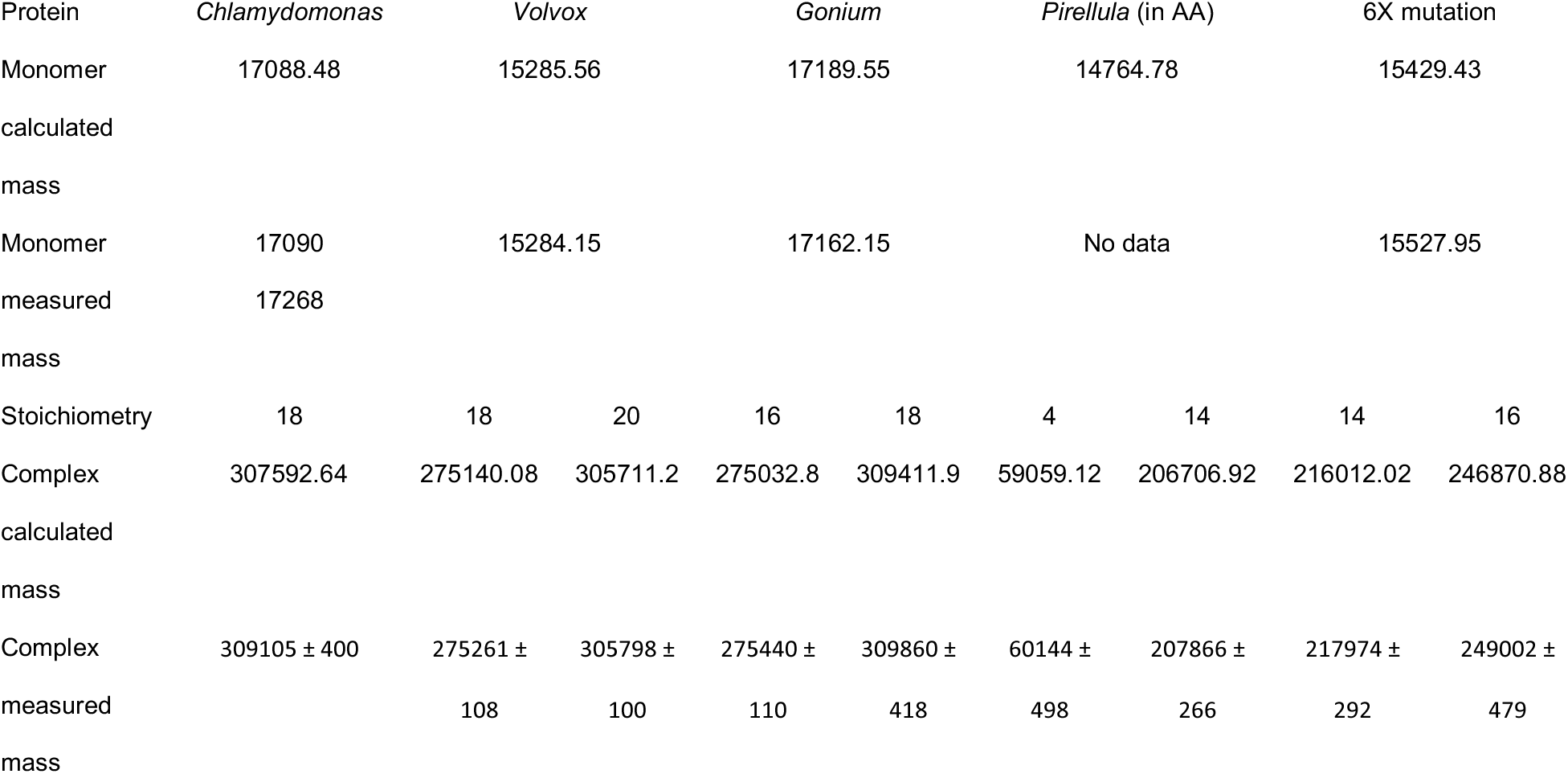
Calculated and measured mass of hub assemblies in this study.

The *Volvox* CaMKII hub domain formed oligomers with molecular weights of 275,261 ± 108 and 305,798 ± 100 Daltons, corresponding to complexes of 18 and 20 subunits, respectively (**Figure 3B**). Oligomerization states were also observed to be concentration independent (spectra acquired at 14 µM and 1.4 µM protomer concentration), suggesting that both the 18-subunit and the 20-subunit complexes are also stable assemblies. At a protomer concentration of 14 µM, the relative populations of the 18-subunit and 20-subunit complexes was roughly 8:1.

The *Gonium* CaMKII hub domain formed oligomers with molecular weights of 275,440 ± 110 and 309,860 ± 418 Daltons, corresponding to complexes of 16 and 18 subunits, respectively (**Figure 3C**). The mass spectrum was acquired at 250 µM protomer concentration. At this concentration, the relative population of 16-subunit to 18-subunit complexes was roughly 5:2. The concentration dependence of the oligomerization states was not tested for this isoform, but given the high sequence identity with the two other algal CaMKII hubs, it is reasonable to assume that the *Gonium* 16-mers and 18-mers are also stable assemblies with defined stoichiometry.

The oligomerization states of the CaMKII hub domain encoded by the bacterium *Pirellula sp. SH-Sr6A* were also assayed by mass spectrometry. This protein is 52% identical in sequence to the human CaMKII-α hub domain, and was expressed and purified in the same manner as the green algae isoforms. The *Pirellula* CaMKII hub forms oligomers with molecular weights of 207,866 ± 266 and 60,144 ± 498 Da, corresponding to complexes of 14 and 4 subunits (**Supplementary Figure 1**). Only the 14-subunit oligomer is predicted to form a closed-ring assembly.

### Structure of the *Chlamydomonas* CaMKII hub domain

We crystallized the hub domain from *Chlamydomonas reinhardtii* and determined the structure 3.0 Å resolution, with 9 subunits in the crystallographic asymmetric unit. An 18-subunit closed ring is generated by a crystallographic two-fold axis of rotational symmetry. The structure of an individual CaMKII hub domain is highly conserved between human and *Chlamydomonas*, with a r.m.s. deviation of 0.9 Å over 115 C_α_ atoms (**Figure 4A**). As in the metazoan CaMKII hub structures, a large N-terminal α-helix is cradled by a highly curved antiparallel β-sheet in the *Chlamydomonas* hub.

**Figure 4.**
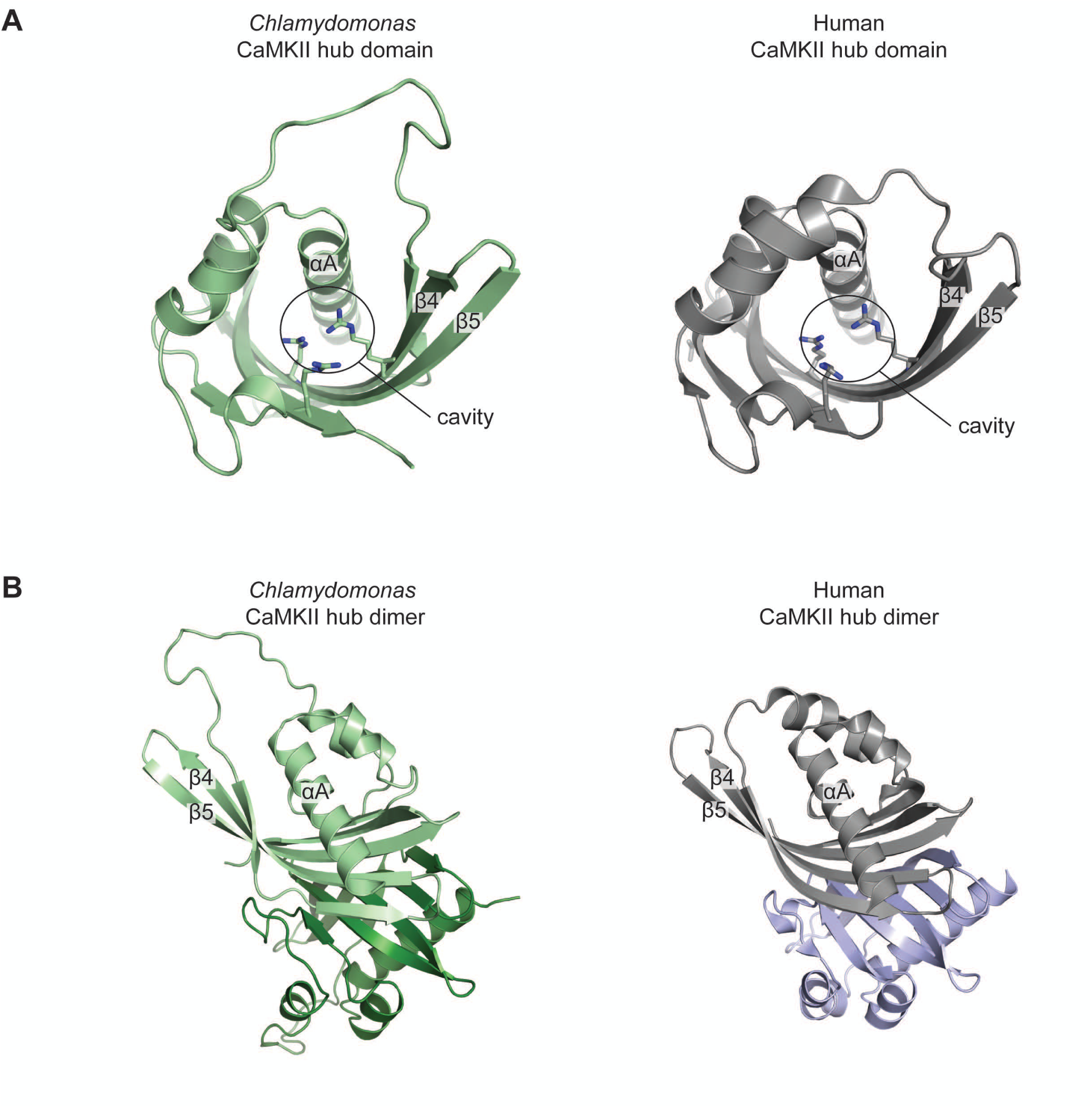
Comparison of the *Chlamydomonas* and human hub domain structures. (A) *Chlamydomonas* and human domains have a highly conserved structure, with a large N-terminal helix cradled by a highly curved antiparallel β-sheet. The conserved arginines in an internal cavity are circled. (B) The structure of the hub domain vertical dimer is highly conserved between the human and *Chlamydomonas* proteins.

The hub domains of human and choanoflagellate CaMKII contain a deep internal cavity containing three conserved arginine residues ^5,7^. The depth of this cavity, and the presence of the three arginine residues within it, distinguishes CaMKII hubs from other proteins with a similar fold, such as ketosteroid isomerases and NTF2. This cavity, and the three arginine residues within it, is conserved in the *Chlamydomonas* hub. In ketosteroid isomerases, the corresponding cavity is the enzymatic active site, but it does not contain the arginine residues. In NTF2, the corresponding cavity is the binding site for a peptide sidechain. The function of this cavity in CaMKII is unknown, but the close similarity in structure and residue composition between the *Chlamydomonas* and metazoan hubs suggests that there may be a binding function associated with this cavity in CaMKII that is conserved.

The structure of the CaMKII hub domain vertical dimer is highly conserved between metazoans and *Chlamydomonas* (**Figure 4B**). In human CaMKII, a striking feature is the presence of 10 histidine residues at the equatorial interface between subunits of the vertical dimer. This feature is conserved in the *Chlamydomonas* hub, which contains 8 histidine residues at the dimeric interface, 6 in the same positions as in metazoan CaMKII hubs. Up to 4 interfacial histidine residues are found in some ketosteroid isomerases and NTF2 isoforms, but the more extensive use of histidines at the interface appears to be a distinctive feature of the CaMKII-like hubs. The importance of these residues for the integrity of human CaMKII is underscored by the recent observation that mutation of one of these residues in human CaMKII-α (His 477) leads to a destabilization of the holoenzyme and is associated with congenital defects in intellectual development ^18^. The unusual utilization of multiple histidine residues at the dimer interface of the CaMKII hubs, largely preserved in the *Chlamydomonas* hub, may point to a pH sensitivity of the stability of the assembly.

Nine vertical dimers are assembled side-by-side in the *Chlamydomonas* hub to form a closed ring assembly (**Figure 5**). The relative arrangement of vertical dimers is the same as that seen in human CaMKII ^5^. A two-fold axis of symmetry in the equatorial plane runs through each vertical dimer interface and is perpendicular to the vertical axis of nine-fold rotational symmetry running through the center of the assembly.

**Figure 5.**
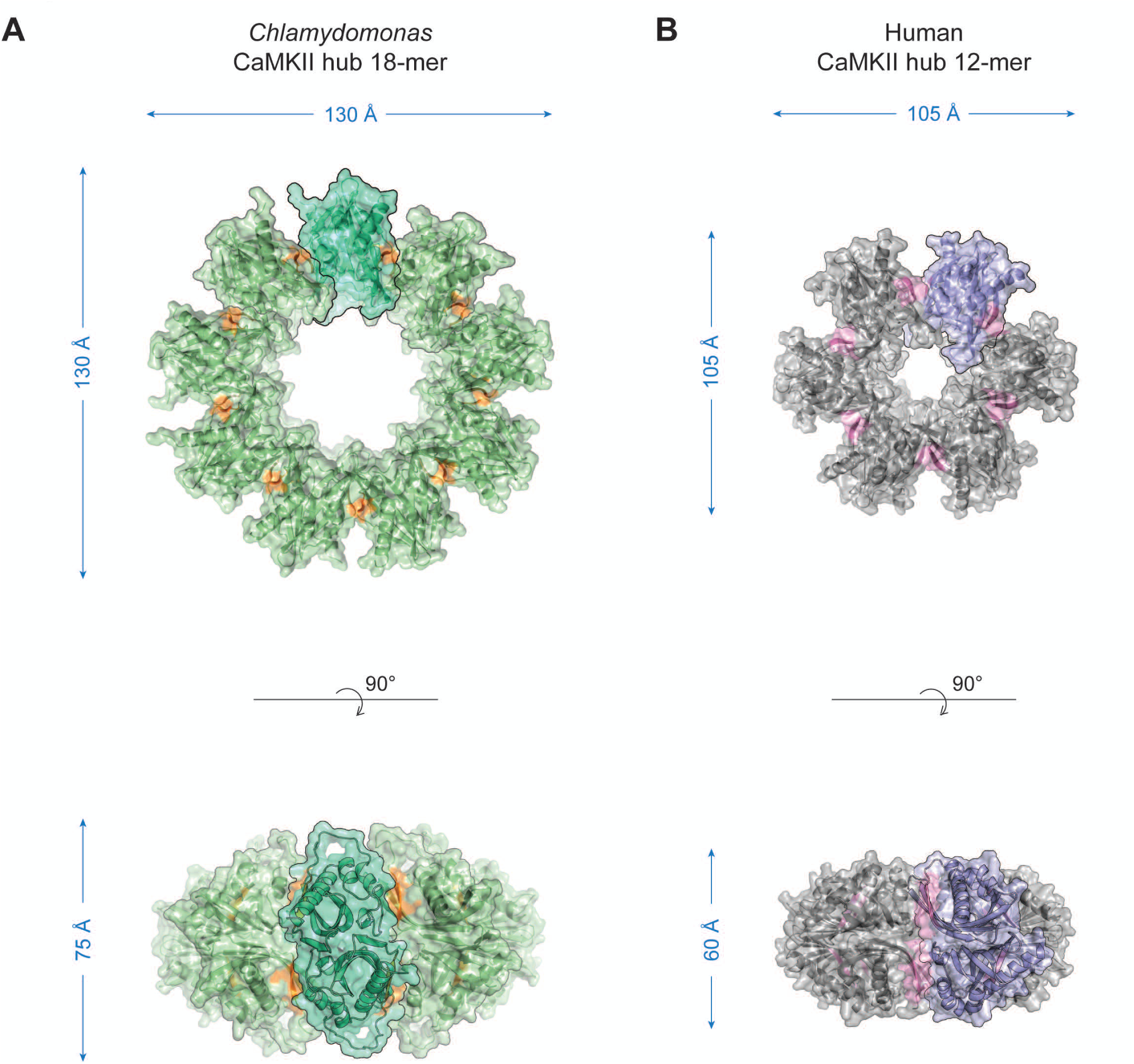
The *Chlamydomonas* CaMKII hub domain assembly. (**A**) The *Chlamydomonas* CaMKII hub domain forms an 18-subunit assembly. The protomers are arranged as nine vertical dimers arranged side-by-side. The upper edge of the β-sheet of each protomer is colored in orange to help delineate the subunits. A vertical dimer of protomers (Stratton et al. 2014) is also highlighted in teal for clarity. (**B**) The dodecameric human CaMKII hub domain assembly is presented for reference. The β-sheet edges are rendered in pink and a vertical dimer is highlighted in purple.

Changes in the twist and curvature of the central β-sheets of hub domains allow different CaMKII hubs to adopt different oligomerization states, while preserving the interfacial interactions that stabilize the assembly ^7^. A lower degree of twist or curvature is seen in the *S. rosetta* CaMKII hub domain, which results in the adoption of an open spiral quaternary structure. An increase in curvature relative to *S. rosetta* causes other CaMKII hubs to form closed-ring dodecamers or tetradecamers, with the β-sheet slightly more curved in tetradecameric forms than in dodecameric forms. The structure of the *Chlamydomonas* hub extends this pattern, with the β-sheet more highly curved in the 18-subunit *Chlamydomonas* assembly than in CaMKII hubs that assemble as open spirals, dodecamers, or tetradecamers (**Figure 6**).

**Figure 6.**
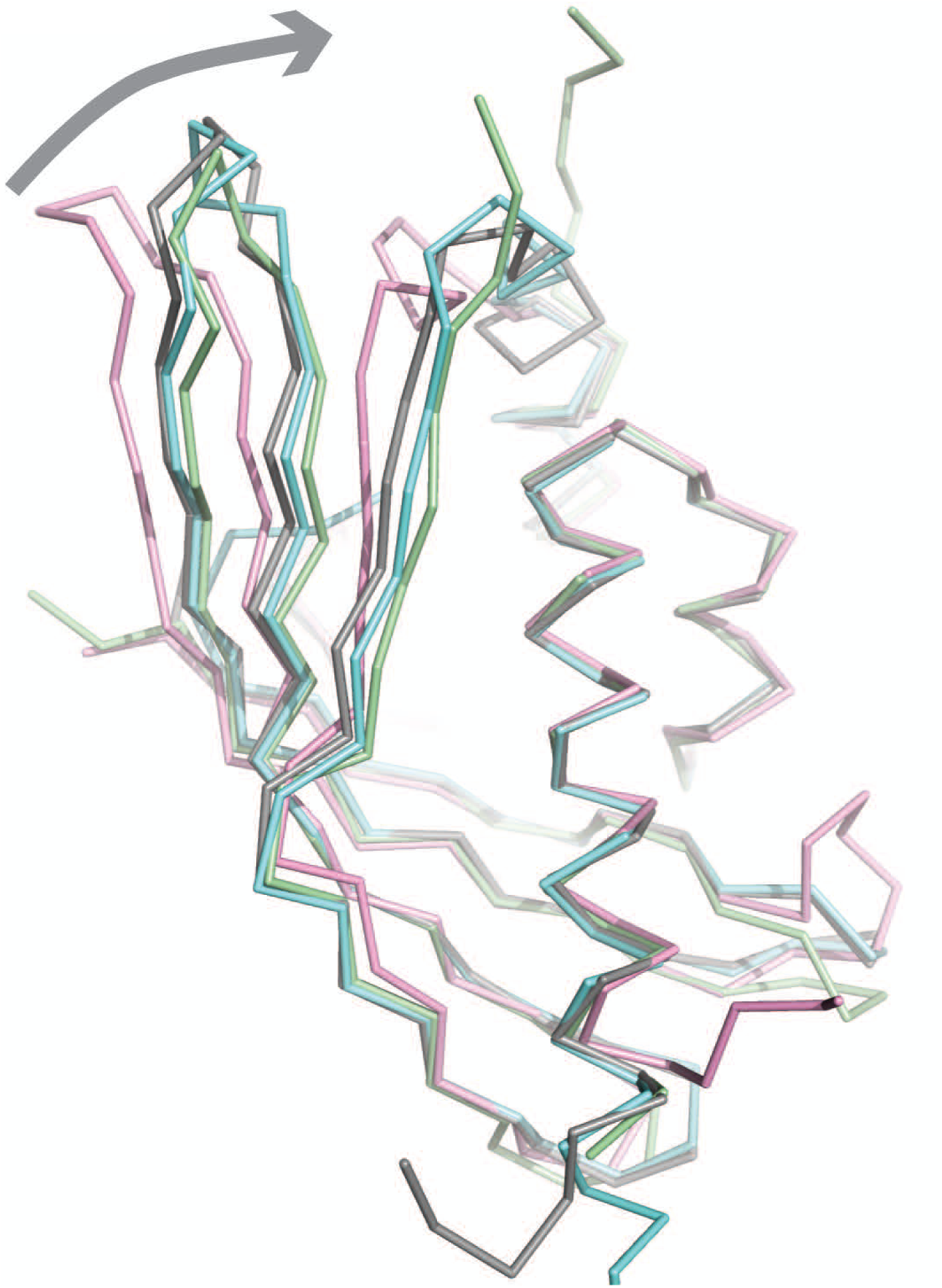
Comparison of the *Chlamydomonas, S.Rosetta*, mouse and human hub domain structures. A single hub domain from each of open spiral *S. Rosetta* (5ig0; pink), human 12-mer (5ig3; grey), mouse 14-mer (1hkx; blue), and *Chlamydamonas* 18-mer (green) were aligned along the central helix and the right most β-sheet section in this view (residues 344-380, 418-430 and 451-469 in the human hub domain). There is a progressive increase in curvature of the β-sheet with assembly size.

### Replacing six residues in the human CaMKII-α hub by the corresponding residues from the *Chlamydomonas* hub increases the number of subunits in the assembly

Four intra-chain hydrogen bonds in the *Chlamydomonas* CaMKII hub domain are not present in the human isoform. These hydrogen bonds appear to stabilize the increased curvature of the β-sheet in the *Chlamydomonas* hub. These hydrogen bonds are predicted to also be present in the *Volvox* and *Gonium* isoforms as well, based on sequence comparison (**Figure 2**).

In the *Chlamydomonas* hub, the sidechain of Gln 12 (residue 355 in human CaMKII-α, UniprotKB Q9UQM7; the corresponding residue numbers for human CaMKII-α are provided in parenthesis in the following discussion), in the middle of the helix αA, forms a hydrogen bond with the backbone carbonyl of Asn 74 (412), on the upper edge of the β-sheet (**Figure 7A**). Gln 12 is replaced by a glutamate in human CaMKII-α, and thus a corresponding hydrogen bond cannot form. Two of the other hydrogen bonds involve conserved arginine residues lining the central cavity. One is formed between the sidechains of Asn 74 (412), on strand β2, and Arg 95 (433), on strand β3 (**Figure 7B**). Asn 74 is replaced by a threonine in the human isoform. Another hydrogen bond is formed between His 125 (464), on strand β6, and Arg 114 (453), on strand β5 (**Figure 7C**). His 125 is replaced by an isoleucine in human CaMKII-α. A hydrogen bond may also be formed between Asn 11 (354), in helix αA, and Tyr 93 (431), on strand β4 (**Figure 7D**). Asn11 is replaced by threonine in human CaMKII-α.

**Figure 7.**
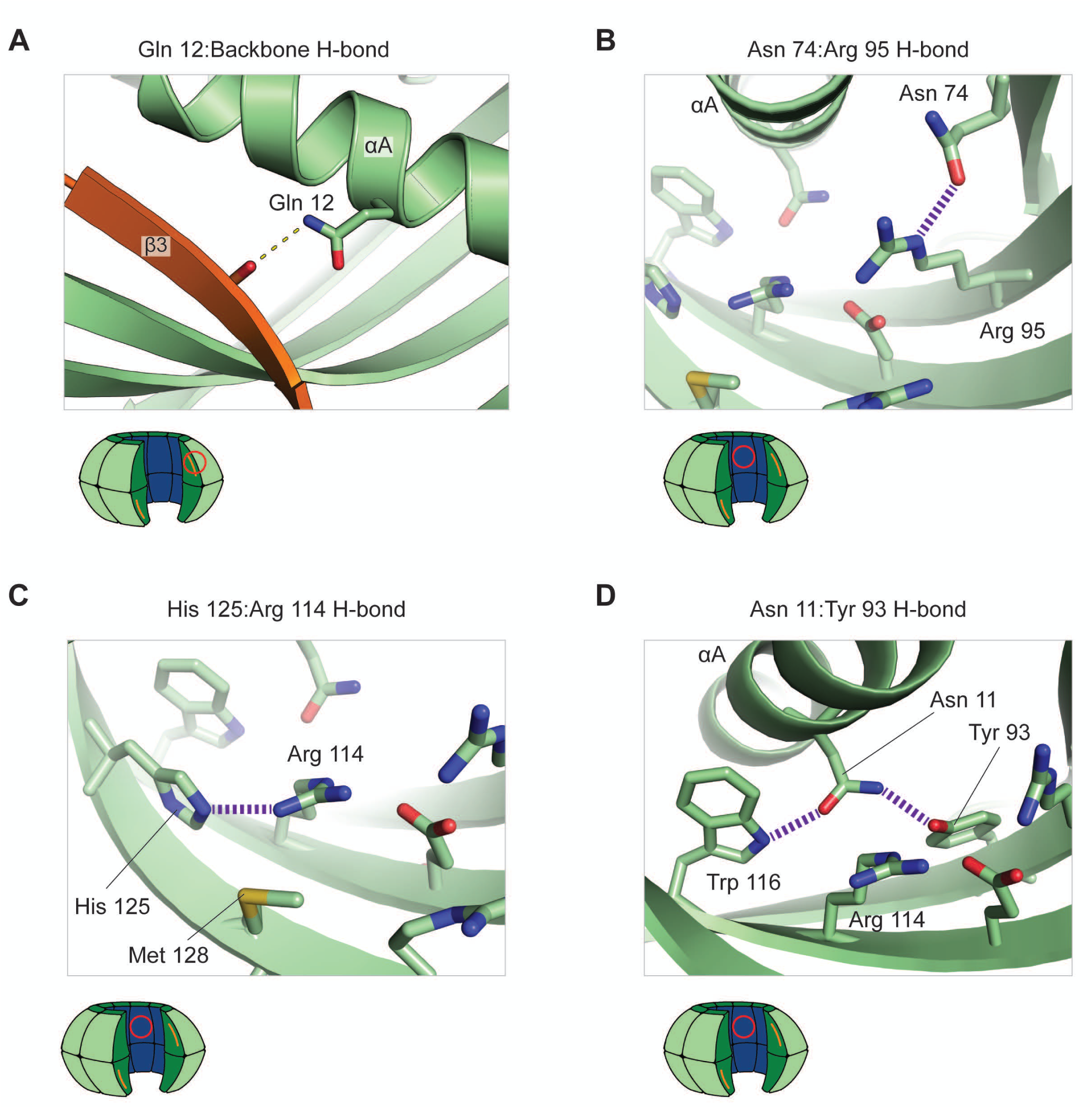
Additional intra-chain hydrogen bonds. (**A**) Glutamine 12 makes an interaction with the backbone carbonyl of Asn 74. The human sequence has a Glutamate at this position which is unable to form the interaction. (**B**) Hydrogen bond between Asn 74 and Arg 92. A threonine residue is present at this location in the human alpha isoform and it is not positioned to form a hydrogen bond with the conserved arginine residue. (**C**) Hydrogen bond between His 125 and Arg 114. The histidine is replaced by an isoleucine in the human isoform. (**D**) Asn 11 forms a hydrogen bond with Trp 116 and also potentially a hydrogen bond with Tyr 93. Asn 11 is replaced by a threonine in the human isoform that forms a hydrogen bond with the tryptophan but cannot simultaneously interact with the tyrosine.

We constructed a mutant form of the human CaMKII-α hub domain with the aim of introducing the four potential hydrogen bonds that are characteristic of the *Chlamydomonas* hub domain. Mutation to the corresponding *Chlamydomonas* residues was performed at four sites in the human hub domain: Thr 354 → Asn, Glu 355 → Gln, Thr 412 → Asn, and Ile 464 → His (human CaMKII-α numbering). We also introduced two additional mutations, Ile 414 → Met and Phe 467 → Met. These methionine residues are conserved across the three algal isoforms, and the structure of the *Chlamydomonas* hub domain suggests that these residues facilitate the increased curvature of the β-sheet. It is important to note that the residues that are mutated are not involved in the lateral protein-protein interfaces between vertical dimers of the hub domain assembly.

The six-mutant CaMKII hub domain was purified and the oligomerization states were assayed by native protein mass spectrometry. The mutant CaMKII-α hub assembles as 14- and 16-subunit assemblies, with a relative population of 7:1. (**Figure 8A**) This is in contrast to results for the wild-type human CaMKII-α hub, which forms roughly equal populations of 12-subunit and 14-subunit assemblies ^7^ and has not been observed to form a 16-subunit assembly (**Figure 8B**).

**Figure 8.**
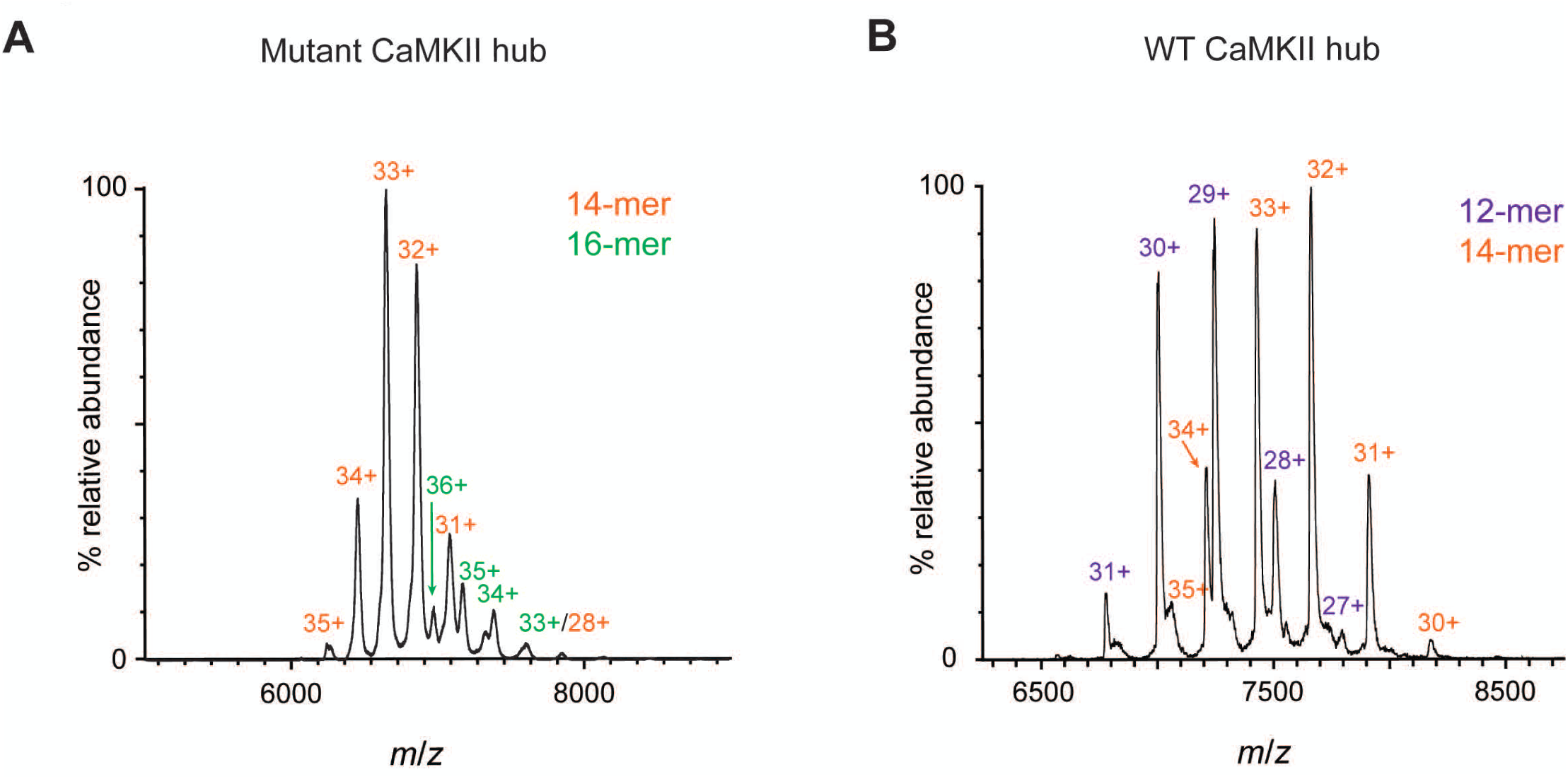
(**A**) Oligomerization states of a mutant human CaMKII-α hub domain. Only 14-mers and 16-mers are formed. (**B**) Oligomerization stats of the WT human CaMKII-α hub domain. Only 12-mers and 14-mers are formed, in roughly equal proportions. Note: 14-mer *m/z* values differ between mutant and WT because the expression tag was not cleaved prior to acquisition of the WT spectrum.

A crystal structure of the mutant CaMKII-α hub domain in tetradecameric form was determined to 2.1 Å resolution (structural overview in **Supplementary Figure 2**). We have not obtained crystals of the 16-subunit form. The asymmetric unit in the crystal contains 7 subunits. In some subunits, Gln 355 (replacing Glu in the wild-type sequence) is positioned to form a hydrogen bond with backbone carbonyl of Asn 412 (**Supplementary Figure 3A**). Asn 412 (replacing Thr 412) and His 464 form hydrogen bonds with their respective partners as predicted (**Supplementary Figure 3B and C**). Asn 354 (replacing Thr 354) appears able to form a hydrogen bond with Tyr 431 but, as in the *Chlamydomonas* structure, the geometry of the bond is not optimal (**Supplementary Figure 3D**).

#### Concluding Remarks

Prior to this work the only known closed-ring CaMKII hub domain assemblies contained 12 or 14 subunits. We now show by mass spectrometry that algal and bacterial proteins with ~50% sequence identity to the human CaMKII-α hub domain form assemblies of 14, 16, 18 and 20 subunits. We determined the structure of a CaMKII hub from *Chlamydomonas reinhardtii* by X-ray crystallography, confirming that it forms a closed ring with 18 subunits.

The structure of the *Chlamydomonas* CaMKII hub domain extends the previously observed correlation between the assembly stoichiometry and the curvature of the central β-sheet. Hub domain flexibility is thought to underlie activation-triggered holoenzyme disassembly in human CaMKII-α ^7,13^. Based on the structure of the Chlamydomonas hub, we introduced six mutations into the human CamKII-α hub. These mutations resulted in a shift in the stoichiometry of the hub, from 12 or 14 subunits in the wild type protein to 14 or 16 subunits in the mutant. It remains to be seen if the mutations that we have introduced will inhibit or in some other way affect activation-triggered holoenzyme disassembly and reassembly in human CaMKII.

## Materials and Methods

### Protein constructs and protein purification

Restriction free cloning was used to insert the relevant CaMKII hub domains into an expression vector harboring a kanamycin resistance gene ^19^. The specific genes from which hub sequences were extracted are: *Chlamydomonas*: UniprotKB A8IHL6, residues 2-134. *Volvox*: UniprotKB D8U3T0, residues 2-137. *Gonium*: UniprotKB A0A150GVZ6, residues 2-136. *Pirellula*: UniprotKB A0A142Y204, residues 2-130. Human: UniprotKB Q9UQM7, residues 345-475. In all cases an N-terminal sequence encoding a hexahistidine tag and Prescission protease cleavage site (MGSSHHHHHHSSGLEVLFQGPHM) was fused to the CaMKII hub gene. The Quikchange protocol was used to insert point mutations into the human hub domain ^20^.

All hub domain isoforms were expressed and purified using a procedure similar to that described for the human CamKII-α hub ^13^. BL-21 DE3 E. coli transformed with the relevant vector were cultured in TB media supplemented with phosphates. Protein expression was induced at ~22°C, OD_600_ = 0.6-0.8 by addition of 1 mM IPTG. Expression proceeded for ~18 hours while shaking at 18°C. All subsequent purification steps were performed at 4°C and all columns were made by GE healthcare.

Cells were pelleted, resuspended in buffer A (25 mM tris, 150 mM KCl, 50 mM imidazole, 0.5 mM DTT, 10% glycerol, pH 8.5 at 4°C) supplemented with DNase and common protease inhibitors, and lysed using a cell disrupter. The soluble lysate was passed over a 5 mL Ni-NTA column and washed with buffer A. Bound protein was eluted in 0.76 M imidazole. A HiPrep 26/10 desalting column was then used to exchange the protein into buffer C (buffer A, but with 10 mM imidazole and 1 mM DTT). Prescission protease was added overnight to remove the expression tags of the *Volvox*, *Pirelulla*, and mutant human hub domains.

Proteins were run over either a superdex-200 or a superose-6 gel filtration column as the final purification step. The column was equilibrated with 25 mM tris, 150 mM KCl, 1 mM DTT, 1 mM TCEP, 5% glycerol, pH 8.0 at 4°C. Fraction purity was assessed by SDS-PAGE. Sufficiently pure fractions were pooled, concentrated, flash-frozen in liquid nitrogen and stored at −80 °C.

### Native protein mass spectrometry

Purified CaMKII hub domain isoforms were exchanged into 1 M ammonium acetate pH 6.9 at 22 °C and diluted to the desired concentrations. PD-25 columns (GE healthcare) were used to perform the buffer exchange. Mass spectra were acquired using a SYNAPT G2-Si mass spectrometer (Waters, Milford; MA, USA). Borosilicate capillaries were pulled to a tip i.d. of ~ 1.5 µm with a P-87 Flaming/Brown micropipetter puller (Sutter Instruments, Novato, CA, USA). Nanoelectrospray ionization was initiated by applying ~1 – 1.5 kV relative to instrument ground on a 0.127 mm platinum wire (Sigma, St. Louis, MO, USA) that was inserted into the borosilicate tips and was in contact with the sample solutions. The instrument was calibrated with cesium iodide (CsI) clusters formed from 20 mg/mL CsI in 70:30 Milli-Q water: 2-propanol solution.

The protein complexes were kept intact using soft instrument interface conditions. The raw data were smoothed with Savitsky-Golay smoothing algorithm (smooth window of mass-to-charge ratio 50 – 100) in Waters MassLynx software. The measured masses of the complexes are slightly higher (up to 1%) than the theoretical masses based on protein sequences due to non-specific adduction of salts ^21,22^.

### Crystallographic analysis

Crystals of the *Chlamydomonas* CaMKII hub domain were grown by the sitting-drop vapor diffusion method. The set drops contained 100 nL of protein solution and 100 nL of well solution. The protein solution was 15 mg/mL Chlamydomonas hub domain in 16 mM tris, 96 mM KCl, 6.4 mM imidazole, 0.64 mM DTT, 6.4% glycerol pH 8.5 at 4°C. The well solution was 0.15 M DL-Malic acid (pH 7.0), 20% (w/v) polyethylene glycol 3350. Drops were equilibrated against 45 µL of well solution and trays were incubated at 20 °C. The cryoprotectant was well solution supplemented with 25% (v/v) glycerol.

Diffraction data were collected at the Advanced Light Source beamline 8.3.1 at 100 °K and wavelength 1.115830 Å. The images were integrated with XDS and then scaled and merged and with Aimless in the CCP4 suite ^23–25^. The structure was solved by molecular replacement with Phenix Phaser ^26^ using two subunits from the tetradecameric human CaMKII-α hub domain assembly (PDB 1HKX) as the search model. The model was refined in coot ^27^. These programs were provided through SBGRID ^28^

Crystals of the mutant human CaMKII-α hub domain were grown by the sitting drop vapor diffusion method. The set drops contained 100 nL of protein solution and 100 nL of well solution. The protein solution was 17 mg/mL mutant hub in 25 mM Tris, 150 mM KCl, 10% (v/v) glycerol, 2 mM DTT, 1 mM TCEP, pH 8.0 at 4 °C. The well solution was 235 mM K_3_•Citrate, 15% (w/v) PEG 3350, pH 8.0 at 20 °C. Drops were equilibrated against 45 µL of well solution. Single crystals appeared overnight, grew to full size within one week, and were harvested and frozen eight days after the conditions were set. The cryoprotectant was 20% (w/v) PEG 3350, 20% (v/v) glycerol, 235 mM K_3_•Citrate pH 8.0 at 20 °C.

Diffraction data was collected at the Advanced Light Source beamline 8.3.1 at 100 °K and wavelength 1.115830 Å. Data was processed as above and phasing was done by molecular replacement with Phenix Phaser ^26^. Subunits from the human tetradecameric CaMKII hub domain assembly (PDB 1HKX) were used as the search model. Model refinement was performed in *coot* ^27^.

## Supplementary material

Crystal structure refinement table. Three supplementary figures.

## Acknowledgements

We thank Moitrayee Bhattacharyya (UC Berkeley) and Margaret Stratton (UMass Amherst) for their advice and for many helpful discussions. We thank James Holton and George Meigs at the Advanced Light Source beamline 8.3.1 for their assistance and guidance during data collection. Supported in part by the National Science Foundation Division of Chemistry under grant number CHE-1609866 (ERW) and by CALSOLV at UC Berkeley.

The authors declare no conflict of interest

