## Supplementary File for "Variation in assembly stoichiometry in non-metazoan homologs of the hub domain of Ca2+/Calmodulin-dependent protein kinase II"

[1]Department of Molecular and Cell Biology, University of California, Berkeley, Berkeley, CA, United States; [2]California Institute for Quantitative Biosciences (QB3), University of California, Berkeley, Berkeley, CA, USA; [3]Howard Hughes Medical Institute, University of California, Berkeley, Berkeley, CA, USA; [4]Department of Chemistry, University of California, Berkeley, Berkeley, CA, USA; [5]Physical Biosciences Division, Lawrence Berkeley National Laboratory, Berkeley, CA, USA

[$]Present address: Intel corporation, Santa Clara, CA, USA

[#]Present address: Department of Chemistry, Columbia University, New York City, NY, USA

[†] To whom correspondence should be addressed:

John Kuriyan

Kuriyan Lab

176 Stanley Hall, MC 3220

University of California

Berkeley, CA 94720-3220, USA

(510) 643-0164

Running title: Discovery and characterization of non-metazoan CaMKII hubs

Supplementary material pages: 8
Tables and figures: 4

Crystal structure refinement table. Three supplementary figures.

**Supplementary Table 1:** Data collection and refinement statistics for *Chlamydomonas* and mutant human CaMKII hub domain assemblies

| | *Chlamydomonas* | Mutant human |
|---|---|---|
| **Data collection** | | |
| Wavelength (Å) | 1.115830 | 1.115830 |
| Space group | P41212 | C 1 2 1 |
| Cell Dimensions | | |
| a, b, c (Å) | 126.46, 126.46, 372.44 | 162.99, 121.34, 56.24 |
| α, β, γ (°) | 90, 90, 90 | 90.00, 108.15, 90.00 |
| Resolution (Å) | 49.04-3.00 (3.08-3.00) * | 47.76-2.10 (2.15-2.10) * |
| Redundancy | 7.3 (7.6) | 3.1 (3.1) |
| Completeness (%) | 100 (100) | 100 (97) |
| $R_{meas}$ (%) | 21.51 (163) | 4 (83) |
| I / σ | 10.0 (1.4) | 13.0 (1.3) |
| CC1/2 (%) | 99 (57) | 99.9 (64.0) |
| **Refinement** | | |
| Rwork / Rfree (%) | 22 / 26 | 21.6 / 25.3 |
| No. observed reflections | 447921 | 187247 |
| No. unique reflections | 61512 | 13211 |
| No. atoms | | |
| Protein | 9235 | 7338 |
| Ligand/ion | - | 18 |
| Water | 11 | 111 |
| R.m.s deviations | | |
| Bond lengths (Å) | 0.012 | 0.009 |
| Bond angles (°) | 1.324 | 0.991 |

*Values in parenthesis correspond to the highest resolution shell

#### SUPPLEMENTARY MATERIALS FIGURE LEGENDS

**Figure S1.** Electrosprayionization mass spectrum (ESI-MS) of the CaMKII hub domain encoded by the bacterium Pirellula sp. SH-Sr6A (UniprotKB AMV33243.1). Two species were detected with molecular weights of 207,866 ± 266 and 60,144 ± 498 Da, corresponding to tetradecamers (14-mers) and tetramers (4-mers). The mass spectrum was obtained at 91 μM subunit concentration.

**Figure S2.** (**A**) Crystal structure of the mutant human CaMKII-α hub domain. The protein crystallized in tetradecameric form. The overall structure is generally the same as the WT tetradecamer (PDB 1HKX), shown in (**B**) as a reference.

**Figure S3**. (**A**) Left panel: Interaction between mutant Gln 355 and backbone carbonyl 412 (blue structure). The glutamine is positioned to form a hydrogen bond with the carbonyl, but the distance is longer than a typical hydrogen bond at 3.6 Å. The WT structure (PDB 1HKX, shown in grey) is aligned on helix α1 for reference. Note that the backbone carbonyl of the mutant structure is closer to the helix, indicating that the β-sheet is slightly more curved relative to WT. Right panel: The glutamine sidechain-backbone carbonyl hydrogen bond in the Chlamydomonas CaMKII hub domain. (**B**) Left panel: Hydrogen bond between mutant Asn 412 and conserved Arg 433 (blue structure). The WT structure (in grey) is again aligned on helix α1 for reference. The WT threonine residue is not positioned to form this hydrogen bond. Right panel: The asparagine-conserved arginine hydrogen bond in the Chlamydomonas structure. (**C**) Left panel: Hydrogen bond between mutant His 464 and conserved Arg 453 (blue structure). Isoleucine is the WT residue at position 464 (grey structure). Right panel: The histidine-conserved arginine hydrogen bond in the Chlamydomonas structure. (**D**) Left panel: Hydrogen bonds between mutant Asn 354 and conserved residues Trp 455 and Tyr 431. In the WT structure (shown in grey) Thr 354 forms a hydrogen bond with the tryptophan but cannot simultaneously interact with the tyrosine. Right panel: Hydrogen bonds formed between Asn 33, Trp 138, and Tyr 115 in the Chlamydomonas CaMKII hub domain.

### Suplementary Figure 1

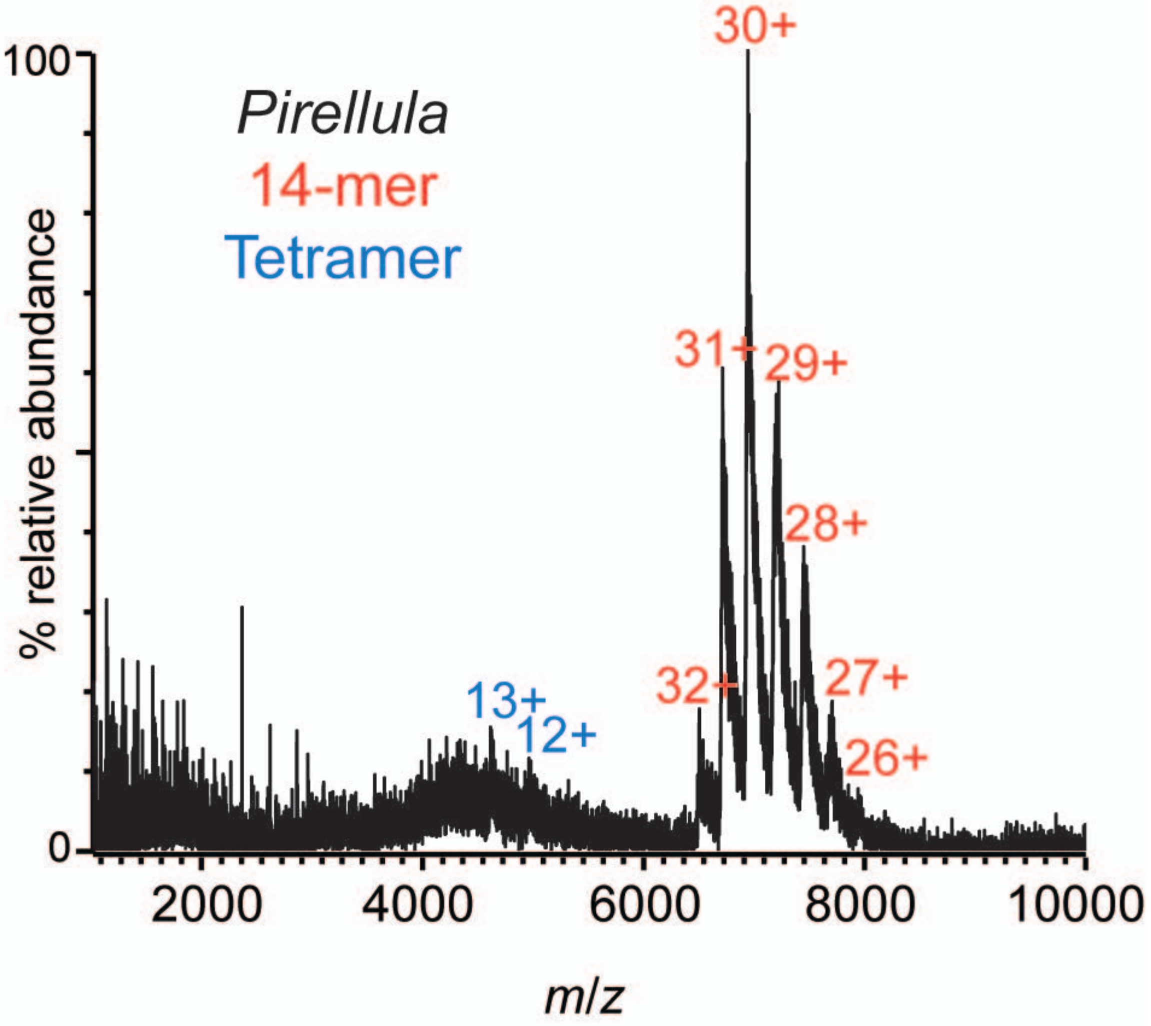

Supplementary Figure 2

**A** Mutant human hub (tetradecamer)

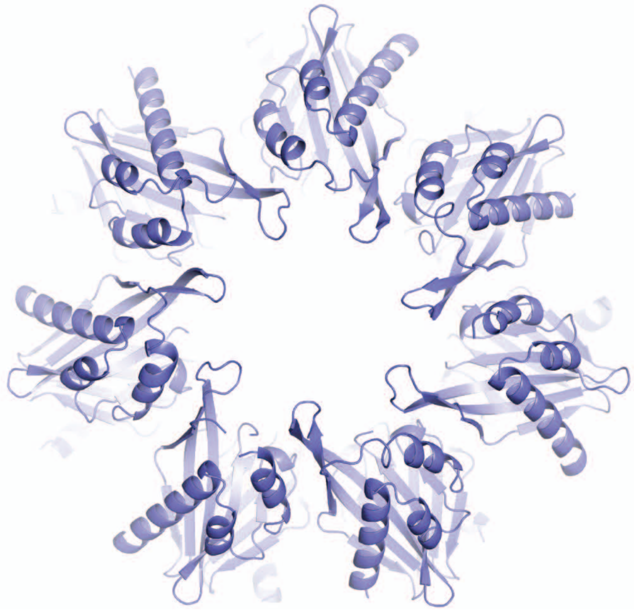

**B** WT human hub (tetradecamer)

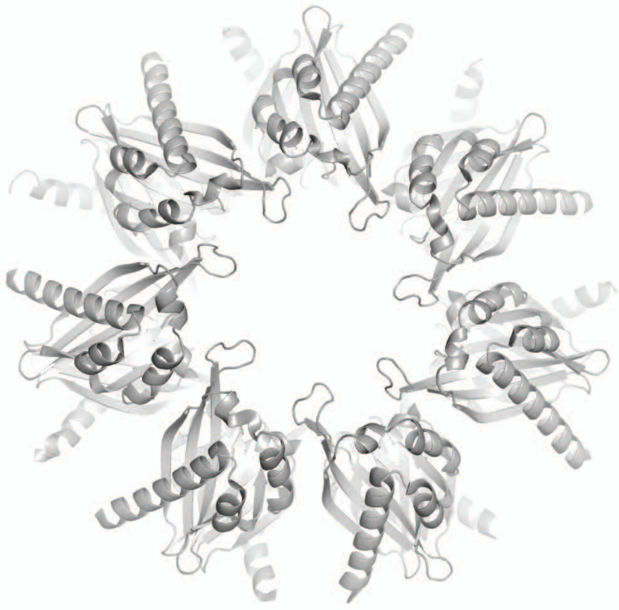

#### Supplementary Figure 3

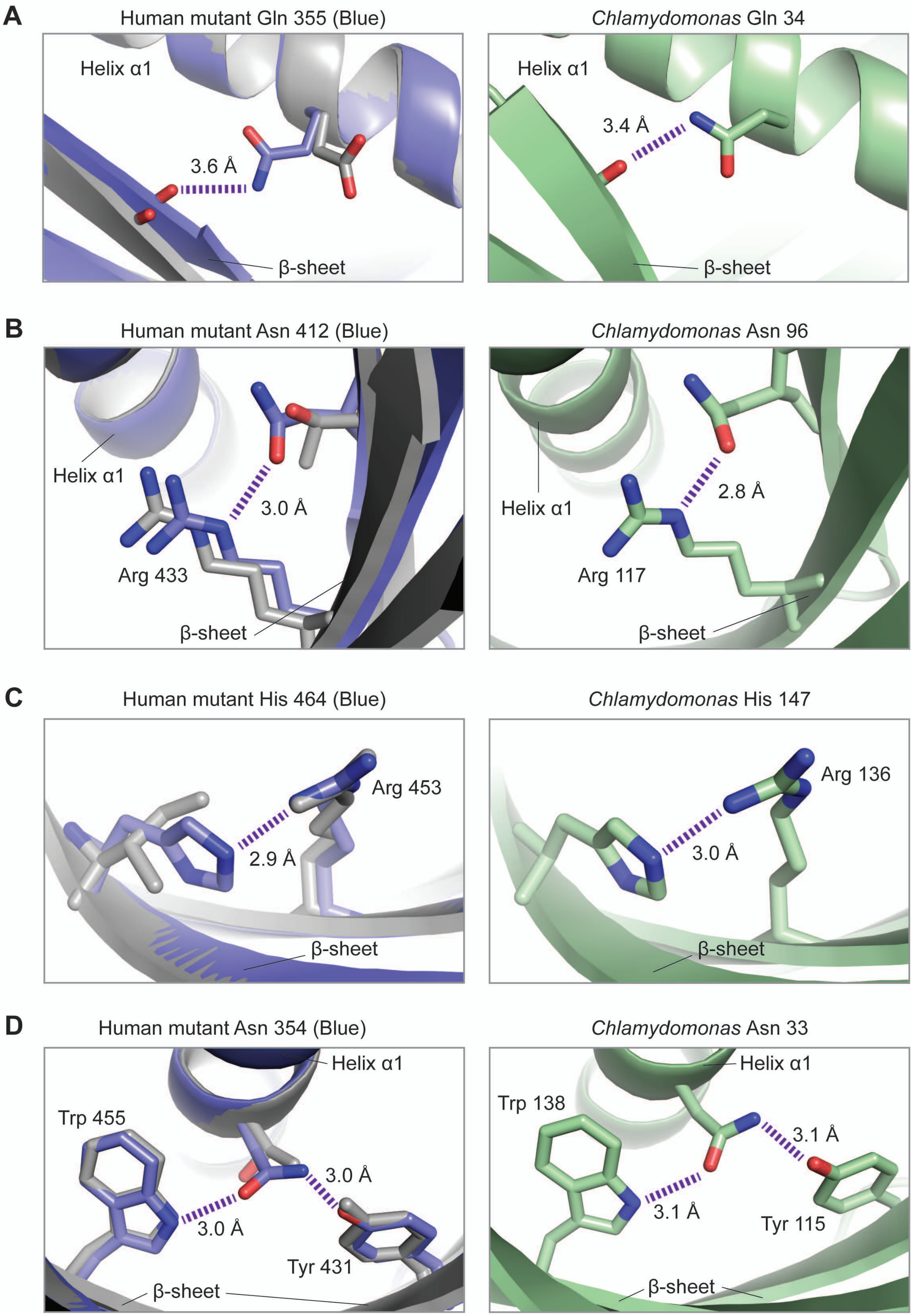
